# Phytochemical evaluation, Embryotoxicity and Teratogenic effects of *Curcuma longa* extract in Zebrafish (*Danio rerio*)

**DOI:** 10.1101/551044

**Authors:** Alafiatayo Akinola Adekoya, Kok-Song Lai, Ahmad Syahida, Maziah Mahmood, Noor Azmi Shaharuddin

**Affiliations:** Department of Biochemistry, Faculty of Biotechnology and Biomolecular Sciences, Universiti Putra Malaysia, 43400 UPM Serdang, Selangor, Malaysia.; Department of Cell and Molecular Biology, Faculty of Biotechnology and Biomolecular Sciences, Universiti Putra Malaysia, 43400 UPM Serdang, Selangor, Malaysia.; Institute of Plantation Studies, Universiti Putra Malaysia, 43400 UPM Serdang, Selangor, Malaysia.; Department of Sciences, College of Science & Technology, Waziri Umaru Federal Polytechnic, Birnin Kebbi, Nigeria.

**Author notes:** E-mail addresses (Alafiatayo, AA); (Syahida, A); (Kok-Song, L); (Maziah, M).

**Keywords:** Toxicology, Curcuma longa, Zebrafish, Flavonoid, FET

## Abstract

*Curcuma longa L.* is a rhizome plant often used as traditional medicinal preparations in Southeast Asia. The dried powder is commonly known as cure-all herbal medicine with a wider spectrum of pharmaceutical activities. In spite of the widely reported therapeutic applications of *C. longa*, research on its safety and teratogenic effects on zebrafish embryos and larvae is still limited. Hence, this research was aimed to assess the toxicity of *C. longa* extract on zebrafish. Using a reflux flask, methanol extract of *C. longa* was extracted and the identification and quantification of total flavonoids were carried out with HPLC. Twelve fertilized embryos were selected to test the embryotoxicity and teratogenicity at different concentration points. The embryos were exposed to the extract in the E3M medium while the control was only exposed to E3M and different developmental endpoints were recorded with the therapeutic index calculated using the ratio of LC50/EC50. *C. longa* extract was detected to be highly rich in flavonoids with catechin, epicatechin and naringenin as the 3 most abundant with concentrations of 3,531.34, 688.70 and 523.83μg/mL respectively. The toxicity effects were discovered to be dose-dependent at dosage above 62.50μg/mL, while at 125.0μg/mL, mortality of embryos was observed and physical body deformities of larvae was recorded among the hatched embryos at higher concentrations. Teratogenic effect of the extract was severe at higher concentrations producing physical body deformities such as kink tail, bend trunk, enlarged yolk sac edema. Finally, the Therapeutic Index (TI) values calculated were approximately same for different concentration points tested. Overall, the result revealed that plants having therapeutic potential could also pose threats when consumed at higher doses especially on the embryos. Therefore, detailed toxicity analysis should be carried out on medicinal plants to ascertain their safety on the embryos and its development.

## 1. Introduction

Plants are source of natural chemical compounds with pharmacological and therapeutic properties. They are widely used for the production of pharmaceutical drugs and play major role in the management of both significant and minor illnesses [1-3]. Although these natural compounds are valuable, some contain toxic compounds with detrimental effect on human’s health [4-6]. Numerous findings have been reported on the toxicity effect of medicinal plants on human organs such as kidneys, liver, heart etc. [7-9] but there are limited reports describing the embryo toxicity and teratogenic effect of *C. longa* extract.

In developing countries, traditional medical practices are the main source of primary healthcare provider. World Health Organization (WHO) reported that 80% of the global population depend on traditional medicine for their healthcare [10]. In recent times the use of natural remedy from plant is becoming more popular among the developed countries as they see medicinal herbs as safe alternatives to orthodox medicines [11].

*Curcuma longa* L. is a rhizome plant that belongs to the family Zingiberaceae, it is often used as traditional medicinal preparations and in everyday culinary. It is a perennial herb widely cultivated in Southeast Asia and distributed throughout world tropical and subtropical regions. The powder form is known as turmeric, popularly used for medicinal purpose and regarded as cure-all herbal medicine with wide spectrum of pharmaceutical activities. Ayurvedic medicine uses turmeric against anorexia, diabetic wounds, biliary disorder, hepatic disorder, and cough while the Chinese traditional medicine claimed it usage for abdominal pains and icterus management [12].

Several therapeutic and pharmacologic properties of *C. longa* have recently been reported; antioxidant activity [13-15], cardiovascular and anti-diabetic effects [16-18] inflammatory and edematic disorders [19-21], anti-cancer [22,23], [26,27], anti-microbial [24-25], hepato-protection [26,27] protection against Alzheimers [28-30], photo protector [12,31]. Although majority of spices and medicinal herbs are commonly presumed to be safe, adverse effects occasionally arise after the consumption of herbal products. The statistical assessment carried out in 2013 by the Malaysian Adverse Drug Reaction Advisory Committee (MADRAC) in conjunction with the National Pharmaceutical Control Bureau and the Health Ministry revealed that 11,473 adverse drug reaction cases were recorded and 0.2% were attributed to herbal medicine [32]. In most nations, toxicity and safety evaluations are not compulsory as a basis for registering herbal product and absence of policies to regulate the production of herbal product contributed to ineffective, sub-standard and possible hazardous consumption.

Despite the widely reported safe pharmaceutical and therapeutical applications of *C. longa*, there is no research finding reporting the embryotoxic and teratogenic effects. Therefore, in this study, the methanol extract of *Curcuma longa* was examined for its flavonoids content, concentration and the *in vivo* embryotoxic and developmental effects using zebrafish embryos and larvae assay as a model.

## 2. Material and Methods

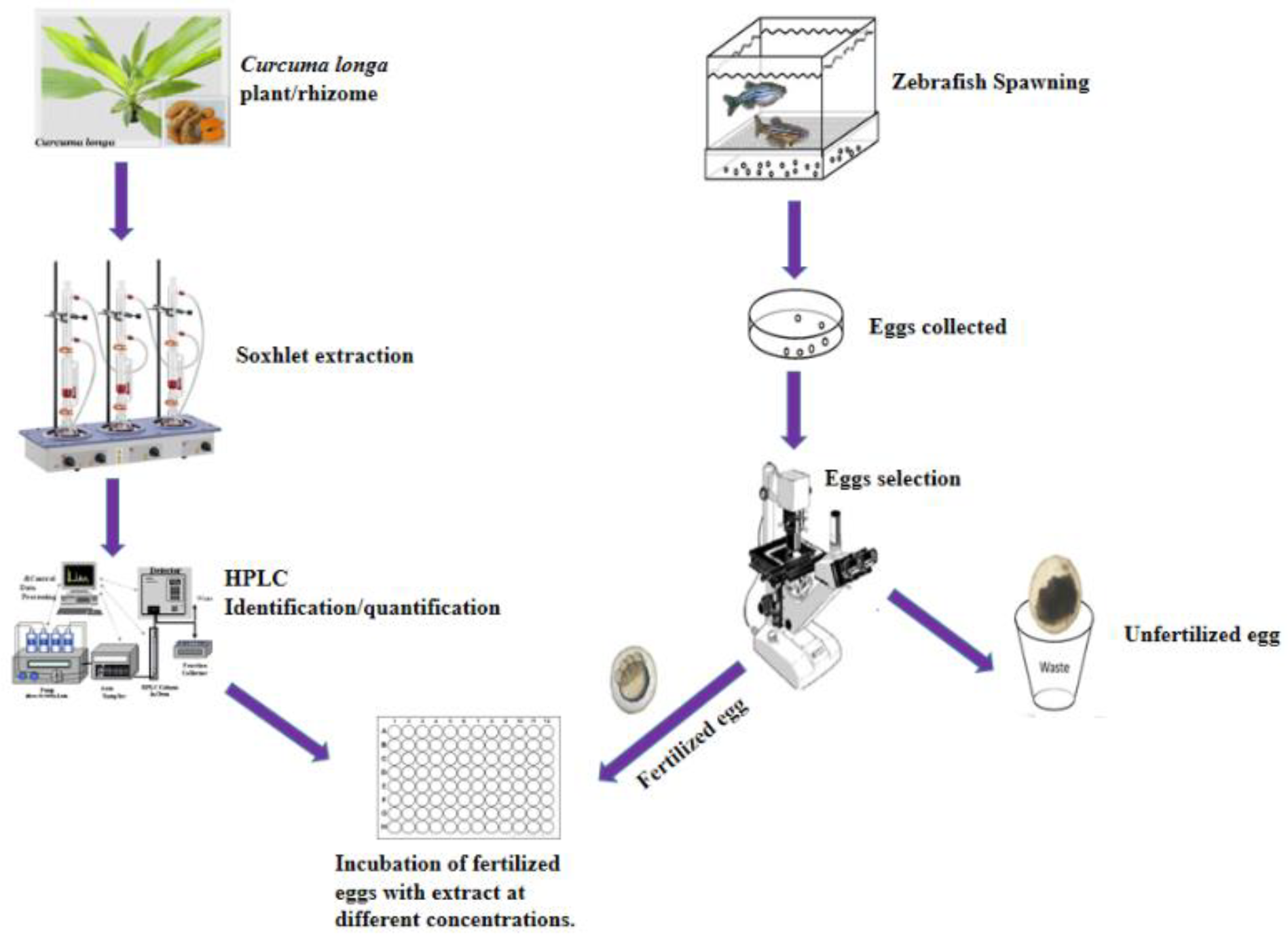
Graphical summary of Method.

### 2.1. Plant Material

The rhizome of *C. longa* was planted at Taman Pertanian Universiti (University Agricultural Park), Universiti Putra Malaysia and the plant harvested after 4 weeks of planting. It identity was confirmed by the residence botanist at the Biodiversity Unit, Institute of Bioscience (IBS), Universiti Putra Malaysia and the plant was deposited in the IBS herbarium with voucher number SK 2849/15 assigned.

### 2.2. Animals and treatment

The maintenance of zebrafish was done in accordance with OECD Fish Embryo Acute Toxicity Test (FET) Draft Guideline of 2006 and approval was given by Universiti Putra Malaysia Institutional Animal Care and Use Committee (UPM/IACUC/AUP No. R024/2014). Adult, wild type, zebrafish (> 6 months old) were bought from a local supplier (Aquatics International Sdn. Bhd. Subang, Shah Alam, Malaysia) kept and maintained at least for 4 weeks for acclimatization to dechlorinated tap water prior to the initial spawning. The adult fish were maintained in 200 L aquarium tank equipped with a continuous flow water system with a maximum density of 1g fish/L tap water at 26±1°C with a constant light cycle of 14:10 hours light-dark at pH 6.8-7.2. They were fed in the morning with brine shrimps (Artemia) supplied by Great Salt Lake Artemia Cysts, Sanders Brine Shrimp Company, Ogden, USA and at noon with dried flakes (TetraMin™ Blacksburg, VA). Nitrate, Nitrite and Ammonia content were checked and maintained below the recommended levels using ammonia test kit. The condition of the fish health was daily monitored.

### 2.3. Production of Fertilized eggs

The wild-type of zebrafish (> 6 months old) with high potential to produce fertilized eggs were selected for spawning. The male and female zebrafish were maintained in aquarium tanks separately with a recommended water volume of 1 litre per fish and a fixed 10 hours of dark periods and 14hrs of light. During the spawning period, excess water filtering and feeding was avoided and cleanness of aquaria and water quality was frequently monitored. Prior to the toxicity testing on embryos, standard method of breeding described in [33] was adopted. Eggs production was from the spawning groups (males and females) at ratio 2:3 respectively. The spawning tank containing 5 L of aquarium water fitted with spawning enhancers which consist of artificial plants and spawn trap (egg collector). 5 spawning tanks of zebrafish were set up to have enough eggs needed for the experiment. Mating occurred within 30 mins in the morning at the time the light was turned on and eggs were collected, the brood zebrafish were subsequently returned back to their resting aquarium tanks. Thereafter, the selection of the fertilized embryos was done and was rinsed trice in embryo medium (E3M) and the fertilized embryos were kept at 280C and allowed to develop for 6 hours.

### 2.4. Curcuma longa extract preparation

*Curcuma longa* rhizomes were harvested after 4 weeks of growth. The collected samples were washed and diced into smaller sizes then dried to a constant weight at 60°C using Memmert Incubators-53L, Model INB 400. The dried sample was ground into a powder form and stored in a clean air tight container.

### 2.5. Extraction

About 0.5 g of each sample was accurately weighed into a reflux flask using GR 200 model of “AND” analytical balance, and 25 mL of 80% methanol was added to each sample. These mixtures were refluxed for 2 hours at 60°C using MTOPS extraction mantle. The resulting mixtures was filtered with Whatman No.1 filter paper, and filtrate was stored in an amber bottle (15 mL) and kept in −20°C refrigerator [34].

### 2.6. HPLC separation and quantification of flavonoid content in Curcuma longa extract

Total flavonoid content extraction for HPLC analysis was carried out by hydrolysis method explained in [35]. About 0.25 g of dried sample was extracted using 10 mL of 60% methanol in aqueous containing 20 mM sodium diethyldithiocarbamate (NaEDTC) as an antioxidant. Subsequently, 2.5 mL of 6 M HCl was then added to the mixture, and this was transferred to a round bottom flask and refluxed for 2hrs at 90°C for the hydrolysis process. The resulting extracts were cooled to room temperature; and then filtered with 0.45 µm filter (Minisart RC15, Sartorius, Germany). Finally, 20 µL was transferred into HPLC vial for the identification and quantification of individual flavonoid using reverse-phase HPLC with 150 × 3.9 mm C_18_ symmetry column.

#### 2.6.1. Preparation of flavonoid standards

The flavonoid standards were prepared for HPLC analysis by weighing 1.0 mg each and dissolving in 1.0 mL of methanol. The standard compounds dissolved were filtered through with 0.45 µm filter (Minisart RC15, Sartorius, Germany). Various concentrations of standard compounds were made to produce standard curve and these were transferred into HPLC vial to quantify the level of individual flavonoid using reverse-phase HPLC with 150 × 3.9 mm C_18_ symmetry column. Similarly, same procedure was done for Apigenin with little modification which is 0.2 mL of dimethyl sulfoxide (DMSO) was used to dissolve 1.0 mg of Apigenin to make a total volume of 1.0 mL. All prepared flavonoid standards were stored in −20°C freezer.

#### 2.6.2. HPLC protocol for flavonoid separation

Using reverse-phase high performance liquid chromatography (HPLC), the rhizome extracts and flavonoid standards were analysed from the Thermo Scientific Ultimate 3000 RSLC System. It comprises of Dionex Rapid Separation Autosampler with NCS-3500RS module with dual – gradient pump and DAD spectral scan. Diode Array detector were used for both UV variants and fluorescence. Separation of compounds was done using reverse-phase separations at ambient temperature using 150×3.9 mm I.D., 4 µm C_18_ Nova-Pak column from Waters (Milford, MA, USA). The mobile phase comprise of 2% acetic acid (aqueous) for solvent A and 0.5% acetic acid (aqueous) plus acetonitrile (50:50 v/v) for solvent B and gradient elution was carried out as follows: 0 - 4 min 2% B, 4 – 40 min 100% B, 40 – 45 min 100% B, 46 – 50 min 50% B. The mobile phase was filtered using 0.45 µm membrane filter under vacuum and column elution was at flow rate of 1 mL/min and detection at a wavelength of 254nm. Flavonoids were identified by comparing retention time and UV spectrum with commercial standards as shown in Figure 1, while the concentration of identified flavonoids were determined using standard curve prepared from commercial flavonoids [35].

**Figure 1.**
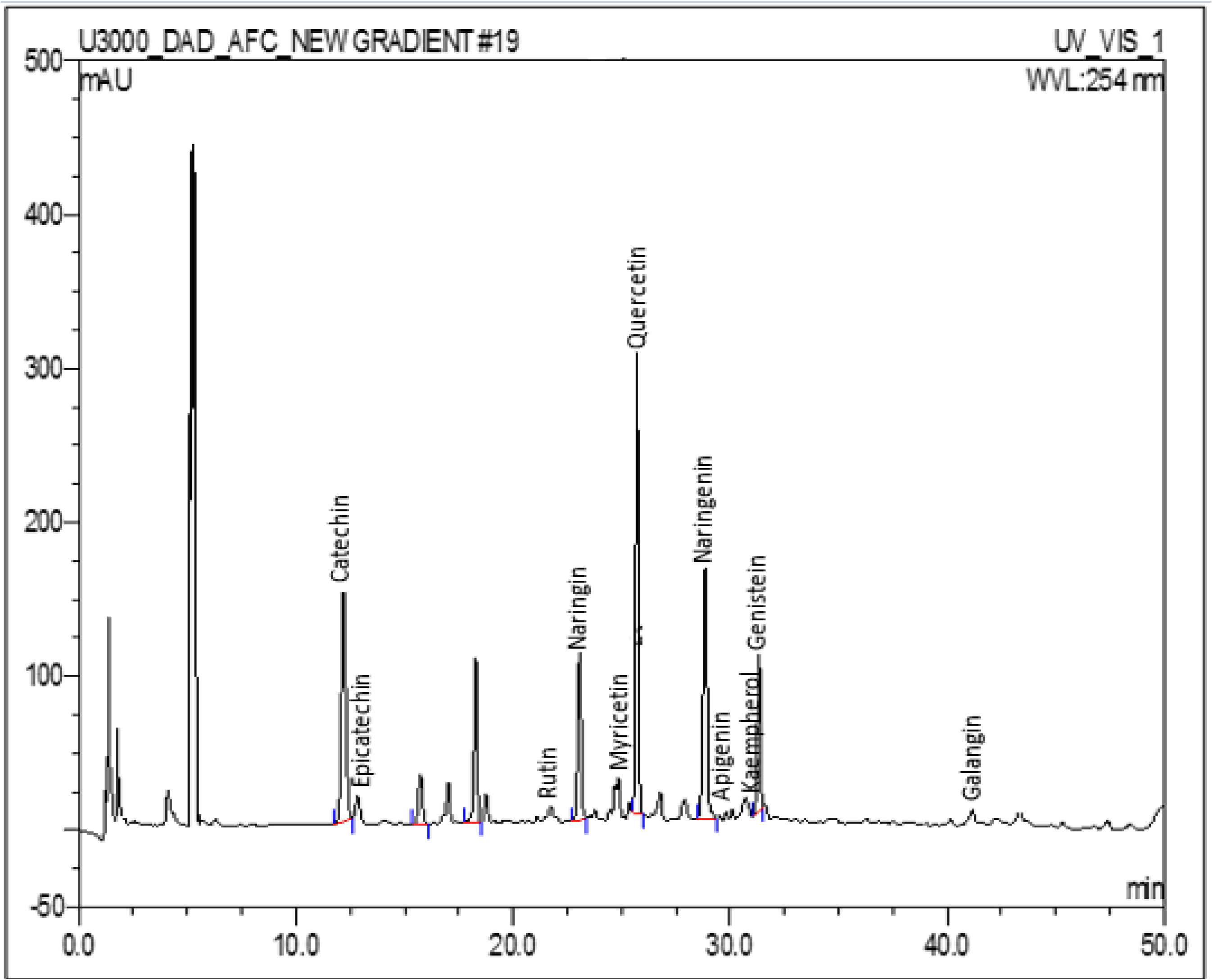
HPLC chromatogram of *Curcuma longa* rhizome extract at wavelength detection of 254 nm and 1.0 mL/min flow rate at different elution time for individual flavonoids detected.

### 2.6. Preparation and dilutions of tests extracts

The extracts (*C. longa*) treatment concentration on zebrafish embryos and larvae were prepared in 24 wells plate separately by diluting 50 µL stock with 49,950 µL of embryo medium (E3M) to produce 125 µg in 0.1% DMSO for *C. longa* and each extract was further diluted 2× dilution factor (1:1) across the well to have 125, 62.5, 31.25, 15.63 and 7.8 µg in 5 mL of E3M having 0.1%. DMSO is the most frequently used solvent for delivery of extracts into zebrafish based assays. In zebrafish embryos and larvae experiment conducted, it was reported that 2.5% concentration of DMSO was well tolerated [36].

### 2.7. C. longa extract Fish embryo acute toxicity (FET) test on Zebrafish

Zebrafish embryos and larvae exposure to the extract were carried out in 24 wells plate according to method described in [OECD. Test No. 236, 2006]. At 6 hours post fertilization (6hpf), selected healthy embryos were washed, examined under the microscope and fertilized embryos were selected for subsequent experiments. 12 fertilized eggs (n=12) at 6hpf for each concentration treatment were treated with the extract of *C. longa* and the experiment was performed in 3 independent replicates in a 24-well plate containing 2 mL of embryo media with 0.1% DMSO containing 125 µg (*C. longa*). It was serially – diluted via 2 folds serial dilution to produce 5 different concentrations of extract *C. longa*. The control (untreated group) was exposed to 5mL of E3M containing only 0.1% DMSO. All the treated groups and control were repeated trice. The development endpoints that were evaluated on both embryos and larvae upon five days exposure were egg coagulation, absent of heart beat in larvae, mortality of embryos and larvae, somites, tail detachment, otolith, eyes and skeletal deformities were recorded each day for five days of exposure (Table 1). The larvae and embryos were subsequently examined with the aid of an inverted microscope (Nikon Eclipse TS 100) to check for malformation of body in each extract concentration for a period of five days exposure. The malformed images of larvae and embryos were captured with Canon Digital camera (power shot A2300 HD). A minimum of 5 different concentrations of *C. longa* were tested for embryotoxic and teratogenic effects on the development of zebrafish embryos and larvae. Among the toxic effects assessed were egg coagulation, hatching and heartbeat while developmental deformities in somites, tail detachment, otolith, blood circulation, heartbeat, motility and skeletal mal-formation were the end of development evaluated for a time period of 5days (120 hours) (Table 1).

**Table 1:**
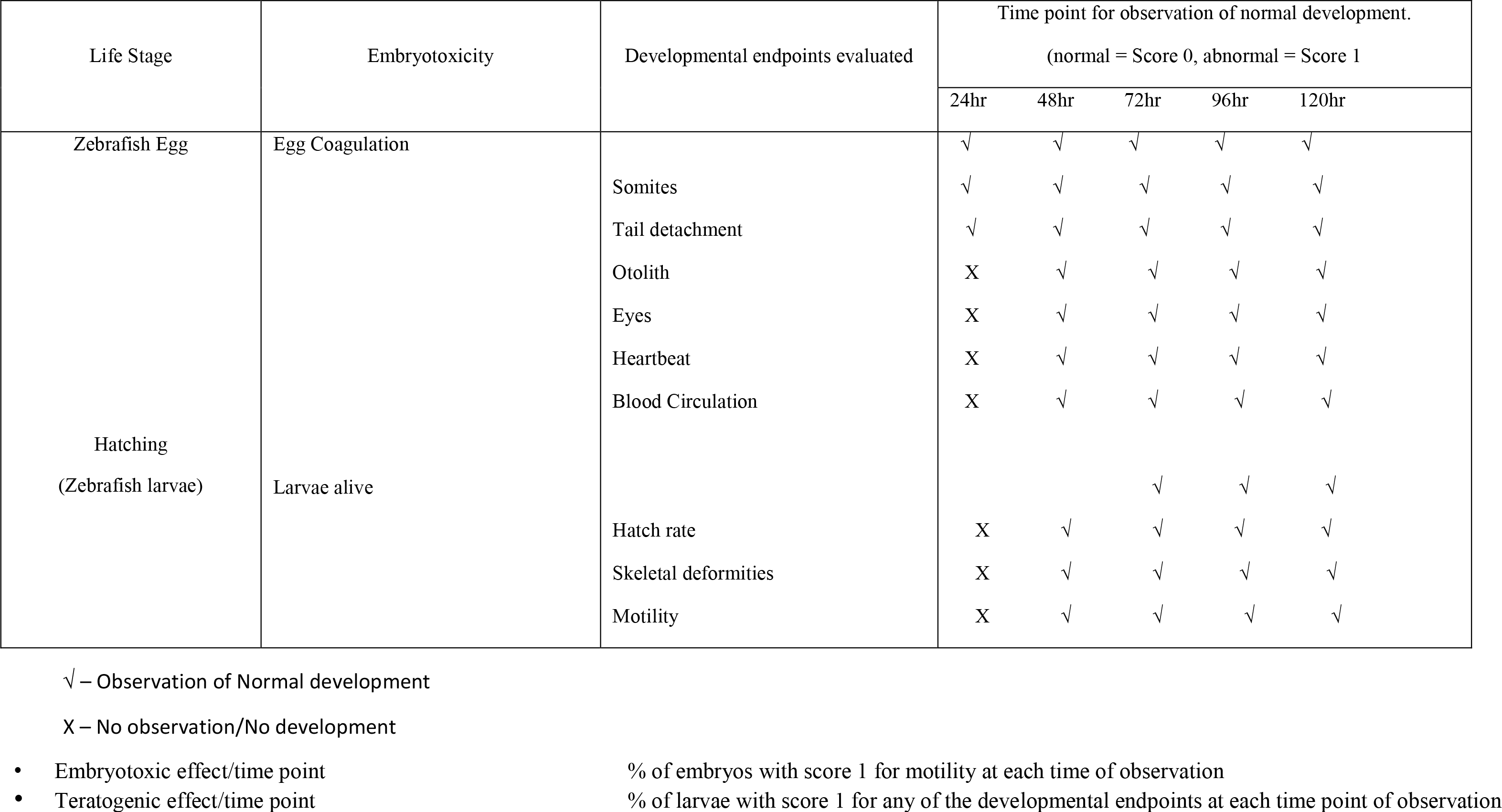
Morphological characteristics evaluated as measures for the teratogenic potency of *C. longa* at different time point.

#### 2.8.1. Evaluations of Zebrafish embryos hatch rate

The zebrafish embryos hatch rate was determined for five days at different concentration of *C. longa* (0 – 125 µg) extracts. The hatching of embryos was taken as the rupture of the chorion for the release of larvae using the inverted microscope.

#### 2.8.2 Evaluations of Zebrafish larvae heart beats

The heartbeat of larvae at five days treatment of *C. longa* extract (0 – 250 µg) was examined in this experiment. The heart beat counting was done by direct visual observation of the zebrafish larval cardiac ventricles using an inverted microscope connected with a computer and camera device. With a stop watch the heart rate was counted per minute.

### 2.9. Therapeutic Index (TI) evaluations

Method described by Selderslaghs et al. (2009) was used for the data evaluation. At time points 24, 48, 72, 96 and 120hpf, mortality/embryotoxicity and morphological changes of the embryos were assessed using inverted microscope (Nikon Eclipse TS 100). Scores were assigned for each characteristic in a binominal manner (‘1’ was assigned for abnormal characteristics and ‘0’ was assigned for normal). Based on the score assigned for particular characteristics, an overall score for percentage effect was created for each treatment in the experiment. An embryo is thought-out to either be normal (all score = 0), malformed or dead for surviving animal (score = 1). In addition, effects were taken as a function of time.

When an increase in mortality is recorded at later time points, malformations incidences were determined as the addition of the incidence at the previous time point for dead larvae/embryos and the incidence for living embryos/larvae at that time. Hence every individual in the experiment was assigned scores for both malformation and mortality at a particular time points. This led to the determination of effective percentage for each concentration at each time point. The embryo toxicity percentage was determined as the ratio of dead embryos and/or larvae over the number of total embryos (12 fertilized eggs) at the exposure start time. Moreover, malformation percentage for 24, 48, 72, 96 and 120 hpf was determined as the ratio of malformed embryos and/or larvae over embryos number that were alive at 24 hpf. Therefore, the resulting output was made up of the cumulative percentage for each time point for observed individual that were dead or malformed.

### 2.10. Dose-response analysis

Using Graph Pad prism, version 5.0, the resulting data from minimum of three independent experiments (n=3) each with twelve (12) replicates (1 embryo per well) per concentration, concentration – response curves for malformed and mortality for each time point was created. The variable shape obtained from the sigmoidal curves adequately fitted the data. The bottom and top curve were set to 0 and 100 respectively with the requisite that percentage near 0 and 100 for effects falls within the concentration range. This concentration – response curve was used in determining the EC_50_ (teratogenic effect) and LC_50_ (lethal/Embryotoxic effects) values. These were derived from four parameter equation describing the curve as follows:

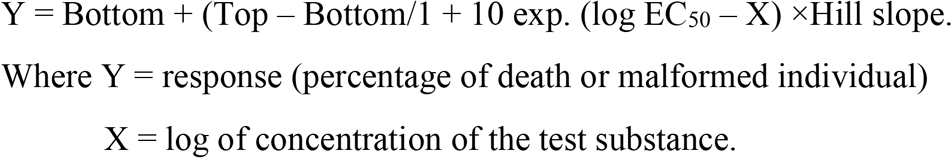

Using calculated LC_50_ and EC_50_ values, a teratogenic index (TI) was calculated as the ratio of LC_50_/EC_50_ for each time point. The higher the TI values, the more the teratogenic effect of the extract tested is specific compare to overall embryotoxicity, as measure by the organism mortality.

### 2.11. Data analysis

Results are presented as mean values ± SEM (n=12) from minimum of 3 independent experiments. With t-test, one way of ANOVA, the statistical significance was determined then Turkey’s post-hoc test was applied using the GraphPad Prism ver.5. Differences were considered significant at p<0.05.

## 3. Results

### 3.1. HPLC Analysis of C. longa extract

The results revealed the presence of certain flavonoids and their concentrations in *C. longa* (Table 2). catechin, epicatechin and naringenin were the three most abundant flavonoid compounds detected with concentrations of 3,531.34, 688.70 and 523.83 µg/mL respectively. Meanwhile, the least three detected flavonoids were naringin, galangin and quercetin with amount concentration of 3.01, 6.54 and 9.43 µg/mL respectively (Table 2).

**Table 2:** Quantified flavonoids in rhizome extracts of *C. longa* using gradient ration of 2% acetic acid (aqueous) to acetonitrile detected at 254 nm.

| Standards<br>Sample | Concentrations (µg/mL) |  |  |  |  |  |  |  |  |  |  |
| --- | --- | --- | --- | --- | --- | --- | --- | --- | --- | --- | --- |
|  | Apigenin | Catechin | Epicatechin | Genistein | Kaempherol | Myricetin | Naringenin | Naringin | Quercetin | Rutin | Galangin |
| *C. longa* | 151.46 | 3531.34 | 688.70 | 63.22 | 101.61 | 76.50 | 523.83 | 3.01 | 9.43 | 112.96 | 6.54 |

### 3.2. Morphological characteristics evaluated as measure for toxicity potency of C. longa extract on Zebrafish larvae and embryos

At 24 post-fertilization (hpf), the embryos were incubated with *C. longa* extract at various concentrations, there was no observable effects at this time point and no hatching of embryo was observed. At 48 hpf, hatching of embryos was observed but no observable effect was noticed in 7.80, 15.63 and 31.25 µg/mL concentrations while at 62.50 µg/mL, bend trunk was observed in some of the group and unhatched darkened embryos were observed in 125.0 µg/mL. At 72hpf, dead hatched larvae were observed at 125.0 µg/mL and morphological deformity such as stunted growth and bend trunk were seen at 62.50 µg/mL concentration. At 96 and 120hpf, kink and bend tail were observed respectively. Dead unhatched embryos were also discovered (Figure 2). For all the concentrations of extract tested, the toxicity effect on each individual was concentration - dependent. Using the percentage of the affected individual (malformation for any observed characteristics) for each concentration, concentration - response curve was produced for each time point (Figure 3). The LC_50_ (for embryotoxic effects/lethality) and EC_50_ (for particular teratogenic effects) data were obtained for the concentration - response curves for all time evaluated based on a minimum of 3 separate experiments (Table 3). The distance between the embryotoxicity and malformation concentration-response curves is taken as a measure of the specific teratogenicity of *C. longa* extract at the time points evaluated. This is also demonstrated by the therapeutic Index (TI) values which are calculated as the ratio of LC_50_/EC_50_ (Table 2). In addition, mortality was observed as a shift to left (lower concentration) as a function of time (Figure 3).

**Table 3:**
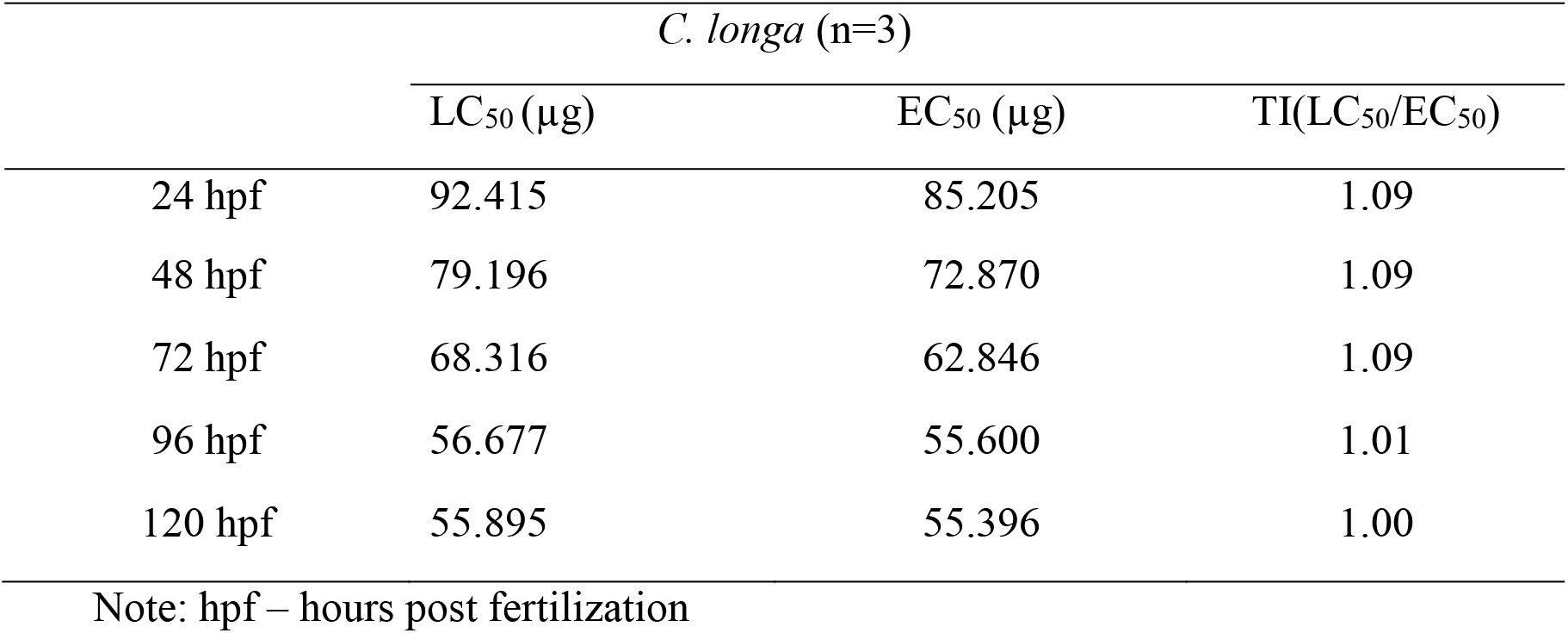
LC_50_, EC_50_ (mean values of 3 independent experiments) and TI values as derived from the concentrations-response curves for *C. longa*.

|  | *C. longa* (n=3) |  |  |
| --- | --- | --- | --- |
|  | LC<sub>50</sub> (μg) | EC<sub>50</sub> (μg) | TI(LC<sub>50</sub>/EC<sub>50</sub>) |
| 24 hpf | 92.415 | 85.205 | 1.09 |
| 48 hpf | 79.196 | 72.870 | 1.09 |
| 72 hpf | 68.316 | 62.846 | 1.09 |
| 96 hpf | 56.677 | 55.600 | 1.01 |
| 120 hpf | 55.895 | 55.396 | 1.00 |

Note: hpf – hours post fertilization

**Figure 2.**
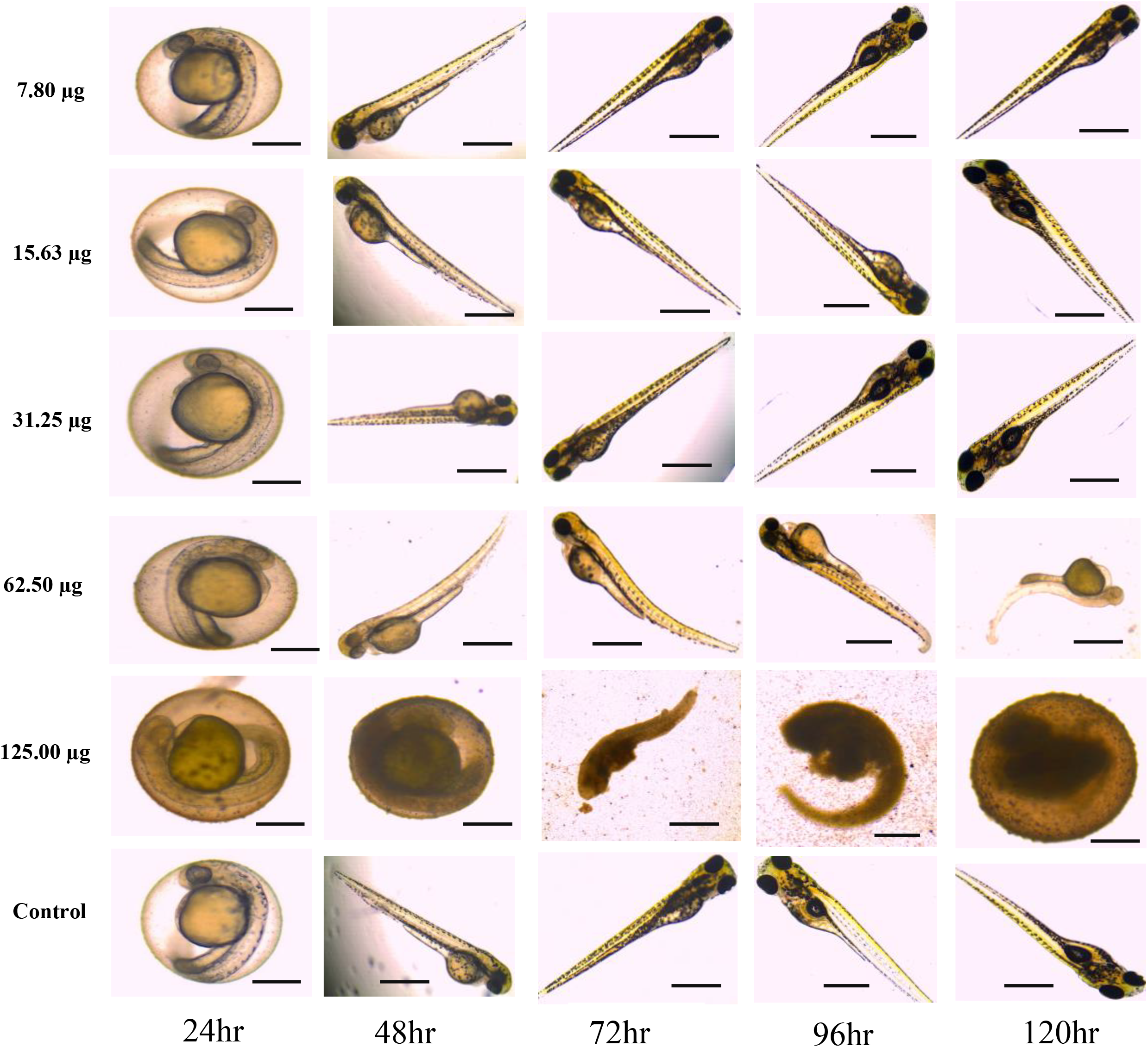
Morphological characteristics assessed as a measure for zebrafish embryotoxicity and teratogenicity of C. longa extract at different time point. At 125 μg the embryos are shown inside the chorion because they died during development and did not reach the stage corresponding to the control.

**Figure 3:**
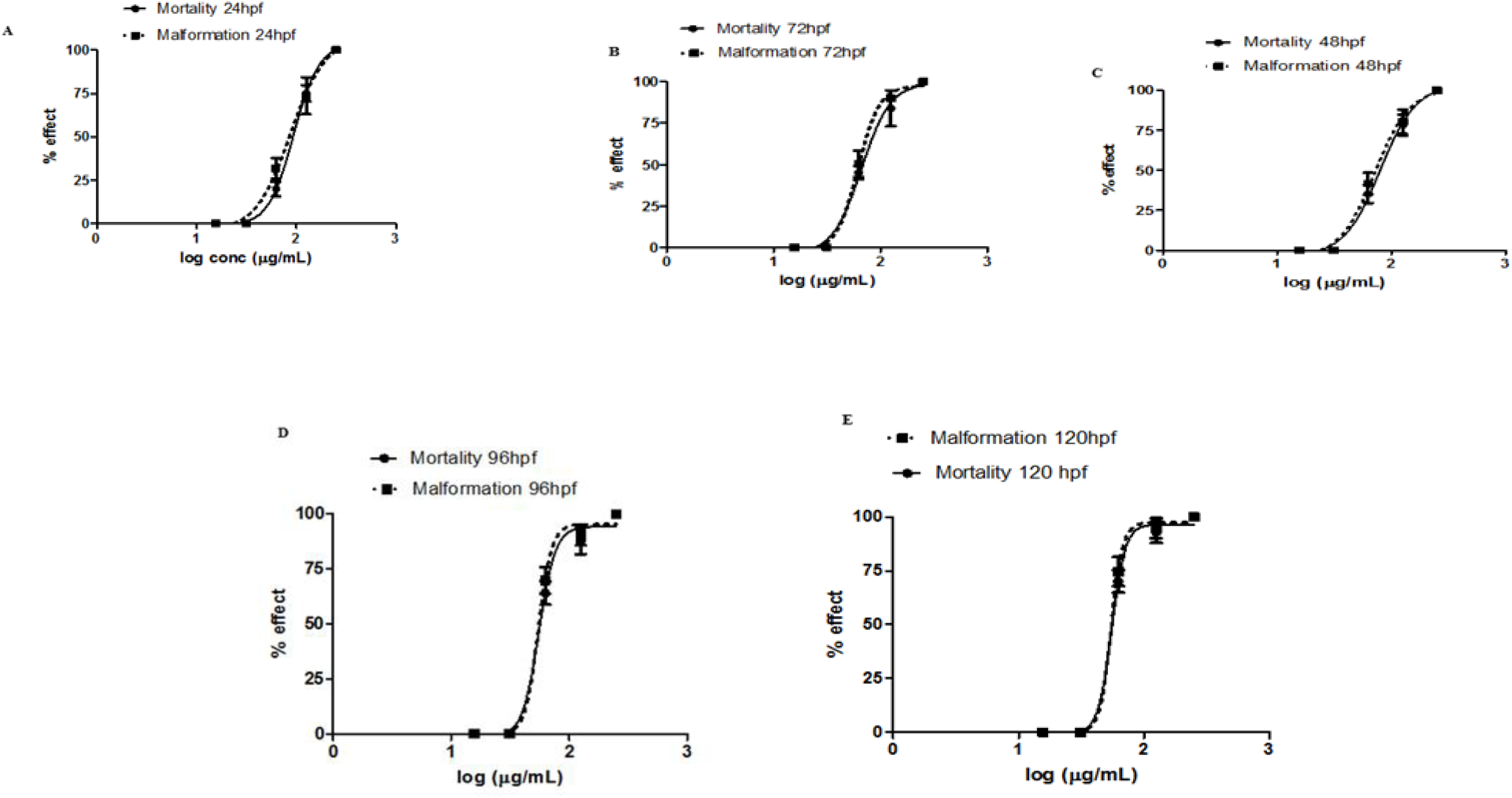
(A-E). Concentration – response curves for malformation and mortality of zebrafish embryos and larvae at different hours of post fertilization (hpf) in different *C. longa* extract concentrations (15.63, 31.25, 62.50, 125.0 and 250.0 µg/mL)

### 3.3. The effects of C. longa extract concentrations on the embryos hatch rate

The hatching rate of zebrafish embryos exposed to varying concentrations of *C. longa* extract displayed delayed hatching at higher concentration of 62.50 µg/mL while no hatching was observed at 125.0 µg/mL as a result of embryos mortality (Figure 2). At 48 hpf, 80% of the embryos were hatched in 15.63 and 31.25 µg/mL concentrations while 100% hatching rate was observed in 7.80 µg/mL which is similar to what was obtained in the control group (Embryos medium – EM) (Figure 4).

**Figure 4:**
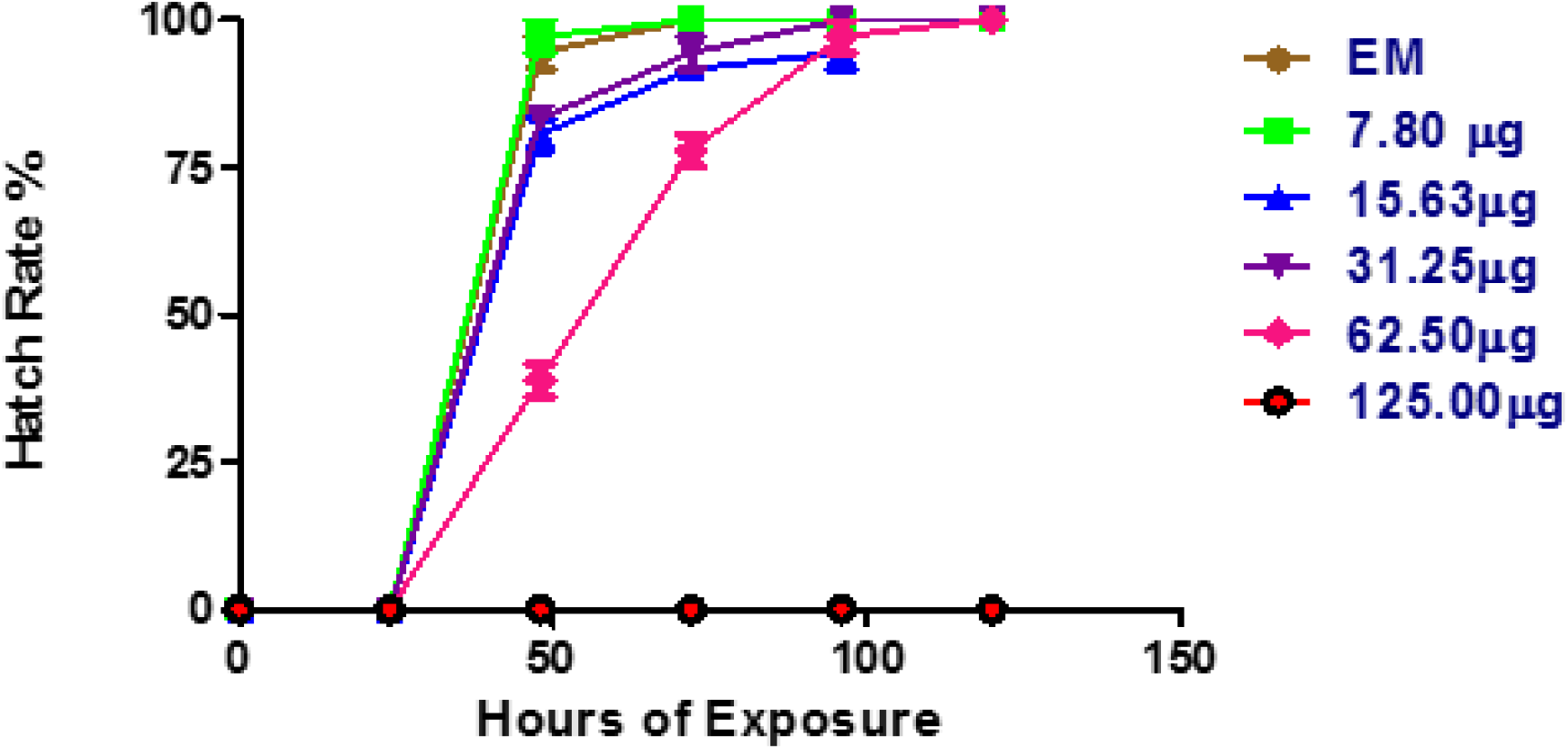
Hatching of zebrafish embryos on exposure to *Curcuma longa* extract.

### 3.4 The effect of C. longa extract on the heartbeat of Zebrafish larvae

The heartbeat of hatched larvae exposed to different concentrations of *C. longa* extract shows no significant difference in the mean heartbeat rate of the control larvae in the concentration range of 7.80, 15.63, 31.25 and 62.50 µg/mL (Figure 5). On the other hand, there was no heartbeat observed at higher concentration range from 125.0 µg/mL above due to mortality of embryos and larvae (Figure 5).

**Figure 5:**
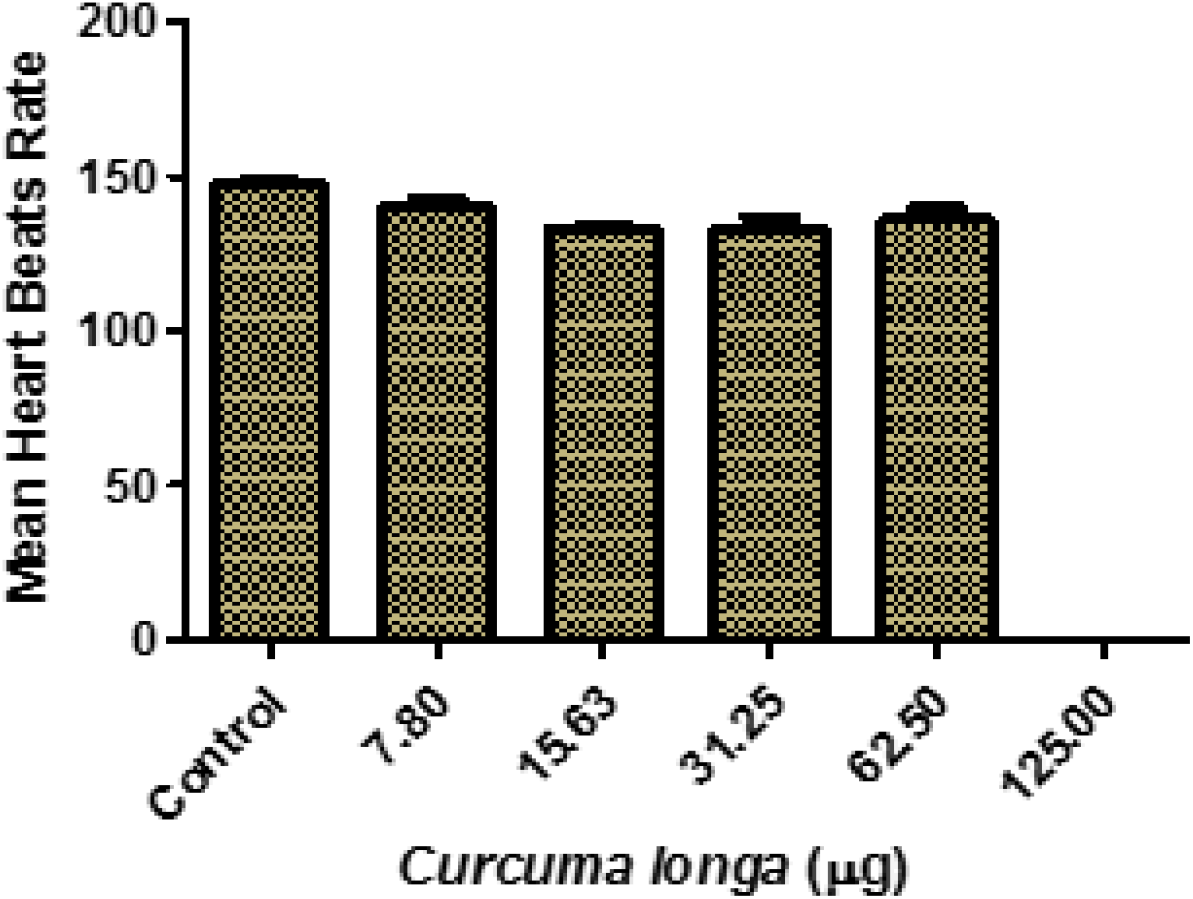
Effects of *Curcuma longa* extract on zebrafish larvae heartbeat. No significant difference at p<0.05 between the control and the tested concentrations group.

## 4. Discussion

Fetal development is a highly organized process in which complex changes are coordinated sequentially in time, and changes at the molecular and cellular levels are integrated to enable manifestation of a particular phenotype in the whole organism. Assessing the embryotoxic and teratogenic toxicity of therapeutic plants on the development of the foetus is important as several products derived from herbal plants claimed to have pharmacological effects are gaining popularity in the global health market without information on their toxicology profile.

This research reveals the detection and concentration of some flavonoids such as Apigenin, Catechin, Epicatechin, Genistein, Kaempherol, Myricetin, Naringenin, Naringin, Quercetin, Rutin and Galangin in the extract of *C. longa* (Table 2) by HPLC analysis. This set of detected flavonoids have been reported to possess some health benefits like antioxidant [38], antifungal and antileishmanial [39]. Kaempherol was reported by Choi et al., [40] to inhibits thrombosis and platelet activation while rutin isolated from *Dendropanax morbifera* L. was also discovered to have anti-thrombotic effect [41]. Furthermore, dietary flavonoids has also been implicated in the amelioration of cataract induced by sugar [42]. Also, the effects of *C. longa* extract on the development of zebrafish embryos and larvae was investigated. The effect of curcumin at different concentrations on zebrafish embryo and larvae had been previously studied by [43] and their findings show a dose-dependent toxic effect of curcumin exposure. At 15 μM of curcumin, all the embryos were reportedly dead within 2days of incubation and all larvae died at 10 μM of curcumin. They further investigated the safety of other polyphenolic compounds such as resveratrol, quercetin and rutin. Unlike curcumin, no toxicity or teratogenic effect was observed in all the polyphenolic compounds tested suggesting that the embryotoxic and teratogenic effects observed in this study may not have been caused by the detected flavonoid compounds in the *C. longa* extract.

However, the findings of Chen et al. [44] reported that synthetic flavonoids such as 7-hydroxyflavone, 6-methoxyflavone, 7-methoxyflavon, 7-aminoflavone and Kaempherol exerted more toxicity on zebrafish larvae as compare to flavone. This suggests that synthetic flavonoids might be toxic at higher concentrations; therefore caution should be taken when consuming them. The toxicity assessment of methanol extract for *C. longa* reveals embryotoxic effect on the zebrafish embryo at higher concentration of 125.0 μg/mL coupled with physiological malformation of larvae development as seen in Fig.2. The deformities were observed to be concentration-dependent (higher concentrations) and increases as the days of exposure increases. The fertilized embryos were exposed to different concentrations of *C. longa* extract ranging from 7.80 ug/mL – 125.0 ug/mL. At 24 hpf no hatching occurred and there was no observable toxicity effect on embryos in all concentrations when compare with the control group. Meanwhile, at 48 hpf hatching of embryo was observed in all concentrations and control group except for 125.0 ug/mL which shows delayed hatching or morbidity of embryos. This suggests possible embryotoxic effect of methanol extract of *C. longa* at higher concentrations. curcumin has been experimentally reported as the most active and abundant compound present in *Curcuma longa* [45]. Dose-dependent toxicity effects of curcumin expose to zebrafish was previously observed and the results showed mortality in embryos at third day of incubation at a concentration of 7.5μM of curcumin and producing deformities in zebrafish larvae [46], which is similar to what is reported in this study. This could possibly be explained that as the exposure of the extract is prolong with increase in days of exposure, there is an increase in the accumulation of the extract until it reaches a concentration that can induce toxicity in the embryos and larvae. The result from this study is similar to the findings of [47], they reported that natural state turmeric exhibited toxicity at higher concentrations on developing embryos.

Also, at higher concentration of 62.50 μg/mL teratogenic effects in the form of deformities in body development was recorded, displaying malformations such as kink tail, bend trunk, physiological curvature and yolk sac edema after 48hpf. The findings of [48] reported slight toxicity at oral consumption and moderate toxicity at intraperitoneal administration of essential oil extracted from oil of *C. longa* cultivated in South western Nigeria in a model mice whereas at lower concentrations of 7.80 ug/mL – 31.25ug/mL no observable malformation was seen (Figure 2) confirming the safety of *C. longa* extract on zebrafish larvae development at lower concentrations [49]. This finding evidenced a significant increase in toxicity effect after hatching at 48hpf resulting into reduction in survival rate, physiological malformation and delayed rates of hatching.

At the concentration of 125 ug/mL (figure 2), an increase in embryo toxicity was observed to be dependent on the time of exposure to extract, and as the time of exposure increases, a decrease in the survival rate of embryo in the chorion was observed. This result suggested that, the accessibility of extract to embryo increases as the time of exposure prolong, leading to the observed toxicity. This could be as a result of weakened or damaged to the embryo protective layer (chorion). In previous experiment conducted by [50], their findings show continuous changes in the protective layer of zebrafish embryo as the age of development advances. They concluded that this could be as a result of the changes in the profile of the chorion protein which might have caused an increase in the opening or widening of the chorion pore channel permitting greater influx of external solute. Furthermore, the effect of the extract on the heartbeat rate of the survive larvae shows no significant difference (Figure 5) when compared with the control and similar result was previously reported by [47], their finding revealed no significance difference in the heart rate of zebrafish embryos treated with raw turmeric and the control. However, the pure curcumin treated embryos demonstrated increase heart rate.

The toxicity effect of curcumin on the proliferation and embryonic development of mouse blastocyst had been previously investigated by [51]. They reported a 7.5-fold higher cell death in curcumin treated blastocysts relative to the control through the generation of ROS and Mitochondria-Dependent apoptotic signaling pathway. On the other hand, recent review on the bioavailability of curcumin and its effects on birth defects was reported by [52]. Their report described the role of curcumin as a scavenger of ROS, which was implicated by [51] as the cause of blastocyst death in mouse. The findings of [52] shows that curcumin can help to ameliorate the toxic effects of certain drugs with teratogenic effects prescribe during pregnancy due to their ability to scavenge ROS.

The concentration – response curve of malformation and mortality of zebrafish embryos and larvae at different time points plotted to obtain the EC_50_ and LC_50_ respectively (Fig 3). The ratio of LC_50_/EC_50_ produces the Therapeutic Index (TI) values for each day of treatment. The TI values is used for ranking the teratogenic effects of any toxic compound i.e. the higher the TI value the greater the teratogenic potential a compound would display [37]. Hence, for this research, the TI values obtained for the 5 days treatment were in close range, suggesting same teratogenic effects throughout the experiment. This is to say, once the deformities in the development are established it cannot be reversed.

## 5. Conclusion

This present research has shown that medicinal herb with potential therapeutic effect could still possess certain toxic effects on embryos and development of larvae especially at higher dosage. Since this extract are usually consumed in their crude form, other phytochemical compounds present in most medicinal plants could subdue the beneficial effect of the extract. Therefore, detailed toxicity assessment should be carried out to establish the safety of extract on embryos and their development as samples confirmed to be safe to organs could still exert toxic effects on the embryo.

## Acknowledgments

We acknowledge the staff of chromatography laboratory, Agro – Biotechnology Institute (ABI), Malaysia for their HPLC technical support. Also, the staff of Taman Pertanian Universiti (University Agricultural Park) and the resident botanist of the Biodiversity Unit, Institute of Bioscience (IBS), Universiti Putra Malaysia.

## Authors’ contributions

All authors contributed equally.

## Abbreviations

DMSO: dimethyl sulfoxide
HPLC: high performance liquid chromatography
OECD: organisation for economic cooperation and development
FET: fish embryo acute toxicity test
hpf: hour of post fertilization

